# Reductive evolution in the structure of the microsporidian proteasome

**DOI:** 10.1101/2022.07.11.499521

**Authors:** Nathan Jespersen, Kai Ehrenbolger, Rahel R. Winiger, Dennis Svedberg, Charles R. Vossbrinck, Jonas Barandun

## Abstract

Proteasomes play an essential role in the life cycle of intracellular pathogens with extracellular stages by ensuring proteostasis in environments with limited resources. In microsporidia, divergent parasites with extraordinarily streamlined genomes, the proteasome complexity and structure are unknown, which limits our understanding of how these unique pathogens adapt and compact essential eukaryotic complexes. We present cryo-electron microscopy structures of the microsporidian 20S and 26S proteasome isolated from dormant or germinated *Vairimorpha necatrix* spores. The presence of distinct densities within the central cavity of the dormant spore proteasome suggests reduced activity in the environmental stage. In contrast, the absence of these densities and the existence of 26S particles post-germination indicates rapid reactivation of proteasomes after host infection. Structual and phylogenetic analyses reveal that microsporidian proteasomes have undergone extreme reductive evolution, lost three regulatory proteins, and compacted nearly every subunit. The highly derived microsporidian proteasome structure presented here reinforces the feasibility of the development of specific inhibitors and provides insight into the unique evolution and biology of these medically and economically important pathogens.

## Introduction

Key biological processes are often executed by essential and complex macromolecular assemblies. Examples include protein synthesis by ribosomes and protein degradation and recycling by proteosomes. Despite the high conservation of these macromolecules in form and function and their universal presence in eukaryotic cells, large variations in assembly, size and complexity can be seen among taxa^1^. Evolution often leads to greater complexity in molecular processes, larger assemblies with novel functionalities, and specialized differences among species. In contrast, for parasitic organisms a reduction in structure and function is often observed. Obligate intracellular parasites frequently undergo this reductive form of evolution through genomic streamlining^2^, a process that is epitomized by microsporidia^3^.

Microsporidia are parasitic eukaryotes with host organisms described from almost all animal phyla^4^. Several species have been isolated from humans and are particularly problematic for individuals with compromised immune systems^5,6^. Microsporidia are notable for many characteristics, such as a lack of innate motility^4^, utility as a biological insecticide^7,8^, and an unusual infection mechanism via injection through a tubular structure^9^. Despite their ecological^10^, biomedical^6^, and agricultural importance^7^, they are perhaps best known for their extreme genome compaction and minimized macromolecular complexes^11–14^. In fact, the microsporidium *Encephalitozoon intestinalis* has the smallest known eukaryotic genome at only 2.3 Mbp^13^. This is half the size of the *E. coli* genome (4.6 Mbp), and 1/65000^th^ the size of the largest confirmed eukaryotic genome (*Paris japonica*, 150 Gbp)^15^.

Microsporidia replicate within a host and have significantly augmented their repertoire of import proteins to facilitate the utilization of host metabolites^16^. Indeed, microsporidia are almost completely reliant on hosts for ATP generation, and the extracellular spore stage of their lifecycle is considered metabolically inert^4^. It is therefore of paramount importance for microsporidian spores to conserve resources, and studies on ribosomes have identified several hibernation or dormancy factors that assist in the inhibition and eventual reactivation of ribosomes post-quiescence^1,12,17^.

Proteasomes play a key role in the conservation of resources by degrading undesirable and misfolded proteins, thereby recycling amino acids. This process requires an estimated 0.25-1.0 ATP per amino acid^18^ and is often inhibited when nutrients are scarce. Interestingly, the mode of inhibition varies based on the limiting nutrient. When nitrogen is unavailable, 80% of proteasomes are degraded by yeast in the first eight hours^19^, despite having a typical half-life in the range of two weeks^20^. On the other hand, when carbon is limiting, proteasomes reversibly accumulate in cytoplasmic puncta known as proteasome storage granules (PSGs)^21^. PSGs are thought to act as a reservoir from which proteasomes can be efficiently recovered once carbon is plentiful^19^.

Eukaryotic 26S proteasomes are macromolecular complexes approximately 2.6 MDa in size, composed of ∼33 different proteins. They can be divided into two sub-complexes: the 20S core particle (CP) and the 19S regulatory particle (RP). While the RP is responsible for substrate recognition, deubiquitination, unfolding, and translocation^22^, the CP houses the proteolytic active sites that hydrolyze targeted proteins. The barrel-shaped CP is arranged in four stacked hetero-heptameric rings, with subunits arranged in a C2-symmetric α1–7β1–7/β1– 7α1–7 conformation^23^. The central cavity formed by these 28 subunits contains six active sites, in β1, β2, and β5 protomers, with caspase-like, trypsin-like, and chymotrypsin-like proteolytic activities, respectively^24^. In the presence of ATP, RPs bind to one or both ends of the CP to form the complete 26S or 30S proteasome^25^. RPs are sub-classified into base and lid complexes. The base contains a ring of six AAA-ATPases (Rpt1-6), three ubiquitin receptors (Rpn1, Rpn10, and Rpn13), and one structural protein (Rpn2)^26^. The lid is typically composed of a deubiquitinase enzyme (Rpn11) and nine structural proteins (Rpn3, 4, 5, 6, 7, 8, 9, 12, and 15), which stabilize the 26S structure during dynamic substrate translocation. Notably, a recent analysis of microsporidian genomes was unable to identify four RP subunits, Rpn3, Rpn12, Rpn13, and Rpn15 (also known as Sem1)^27^, despite high architectural conservation in eukaryotic proteasomes, highlighting the highly derived nature of microsporidian proteasomes. This is of particular interest as proteasomes are tempting targets for selective inhibition, and very few treatments are currently available for microsporidiosis^27^.

In this work, we study the impact of reductive evolution on 20S and 26S proteasome structures using cryo-electron microscopy (cryo-EM). We present the first two structures of microsporidian 20S proteasomes, isolated from either dormant or germinated spores of *Vairimorpha necatrix*, a generalist parasite of lepidopterans. All active sites of the proteasome from dormant spores are occupied by the peptide fragments in conformations incompatible with active digestion. The high occupancy of these fragments and their absence in proteasomes extracted from germinated spores suggest that microsporidia reduce proteasome activity in the extracellular stage and can efficiently reactivate proteasomes upon germination and infection of the host. The existence of a small number of singly- and doubly-capped proteasomes in the germinated sample allowed us to generate an architectural model of the 26S proteasome. Our data are consistent with a highly modified 26S structure missing Rpn12, Rpn13, and Rpn15, and containing a notably truncated Rpn3.

A comparison of yeast and microsporidian proteasomes demonstrates that, while microsporidia have retained the expanded repertoire of 20S subunits characteristic of eukaryotic proteasomes, they have eliminated many of the insertions which distinguish individual α and β subunits. These deletions and truncations have resulted in small-scale structural rearrangements throughout the proteasome, most importantly at the CP gate and in several regions interfacing with the RP. Although active site residues responsible for proteolysis are well conserved, the specificity pocket of the *V. necatrix* β5 subunit adopts an open conformation associated with selective inhibition by several α’, β’ epoxyketone compounds, highlighting the utility of these compounds as possible therapeutics. Finally, a phylogenetic analysis using proteasomal subunits reinforces previous taxonomic groupings and emphasizes the structural streamlining underpinning reductive evolution in microsporidian macromolecular complexes.

## Results

### Isolation and structural characterization of microsporidian proteasomes

The study of microsporidian macromolecular complexes is complicated by a lack of genetic tools to tag or modify proteins of interest and an inability to culture microsporidia in a non-cellular growth medium, leading to limitations in starting material and requiring the use of traditional enrichment procedures. Additionally, to obtain a molecular understanding of the *V. necatrix* proteasome, it is important to study it in conditions consistent with both dormant and activated environments (**Fig. 1a**).

**Figure 1.**
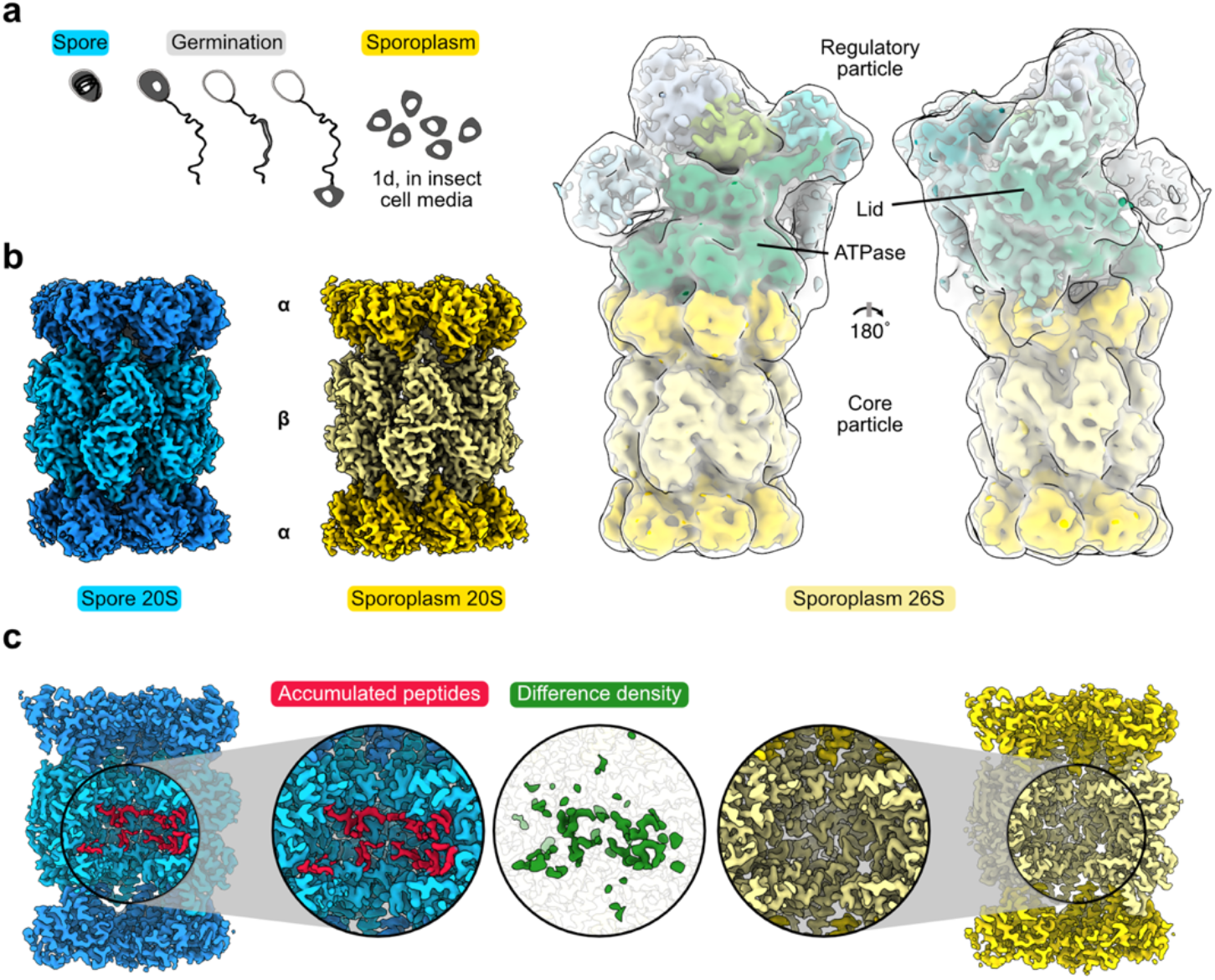
The cryo-EM structure of the microsporidian proteasome, isolated from spores or sporoplasms. **a** Schematic overview of the germination process of a microsporidian spore proceeding from the environmental stage to the release of the sporoplasm through the polar tube. The stages from which proteasomes were isolated are labeled in blue (spore stage) and yellow (sporoplasm stage). **b** Cryo-EM density of spore-isolated (blue) and sporoplasm-isolated (yellow) proteasome reconstructions. The 8.3 Å 26S cryo-EM density is shown with RP subunits colored shades of green, enclosed in a transparent white 16-Å lowpass-filtered map. **c** Slab-view of the full proteasome and zoom-in sections of the central proteolytic cavities. Additional peptide-like density (red) is present in the spore (left) but absent in the sporoplasm isolated proteasome (right). The central circular inlet shows the difference density (green) between 4-Å lowpass-filtered cryo-EM maps from spores and sporoplasm proteasomes.

Although previous work has noted difficulties in matching proteasomal orthologs in microsporidia due to low sequence identities^27^, the resolution of our data allowed us to unambiguously identify all CP subunits (**Supplementary Table 2**). Similar to typical eukaryotes, microsporidian proteasomes are composed of 28 proteins in a C2-symmetric α1–7β1–7β1–7α1–7 arrangement. Most regions involved in essential functions, such as the catalytically active residues and substrate recognition pockets, are structurally well-conserved. On the other hand, many short loops and disordered tails are absent. Additionally, within the central cavity of the CP barrel, we identified peptide-like densities that span multiple β subunits and block the active sites of β1, β2, and β5 (**Fig. 1c**). The identities of the fragments could not be unambiguously assigned using density-informed motif searches in combination with mass spectrometry (**Supplementary Data 1**).

To investigate proteasomes from an active microsporidian stage and determine if these unidentified densities are spore specific, we activated 10 mg of microsporidian spores through germination using alkaline priming^29^. We then incubated the germinated spores for one day in an ATP-rich insect cell medium, gently lysed the fragile sporoplasms via sonication, and purified proteasomes in the presence of ATP using differential centrifugation. Both 20S and 26S particles were clearly visible in these samples via cryo-EM (**Supplementary Fig. 1**), indicating that either the germination or the presence of ATP in the buffer facilitated the association of the RP and CP. The presence of 20S, 26S, and 30S proteasomes allowed us to obtain a volume of the CP at a resolution of 3.2 Å (17,942 particles), as well as a low-resolution volume (∼8.3 Å; 6,442 particles) of the 26S proteasome (**Fig. 1b**). The overall structure of the spore and sporoplasm isolated 20S proteasomes are essentially identical. Both proteasome pores are closed, and the catalytic chamber is not accessible. In contrast to the overall structural similarities, the sporoplasm core particle lacks the peptide-like densities found in the spore 20S.

### Proteasomes from dormant spores accumulate peptide fragments

Chambered proteases house multiple active sites responsible for proteolysis within barrel-shaped cavities^30^. They are found in all three domains of life and are typically utilized as broad-spectrum proteases to digest undesirable proteins into small fragments^30^. Proteasome CPs contain three connected cavities through which unfolded proteins pass: two antechambers created by the α*-*rings, and a catalytic chamber housing the active sites of the β-rings (**Fig. 1b, c, and Fig. 2**). Previous data led to the speculation that the three chambers of the proteasome can act like the multiple stomachs of ruminant animals^30^, allowing for multiple rounds of digestion of peptide fragments.

**Figure 2.**
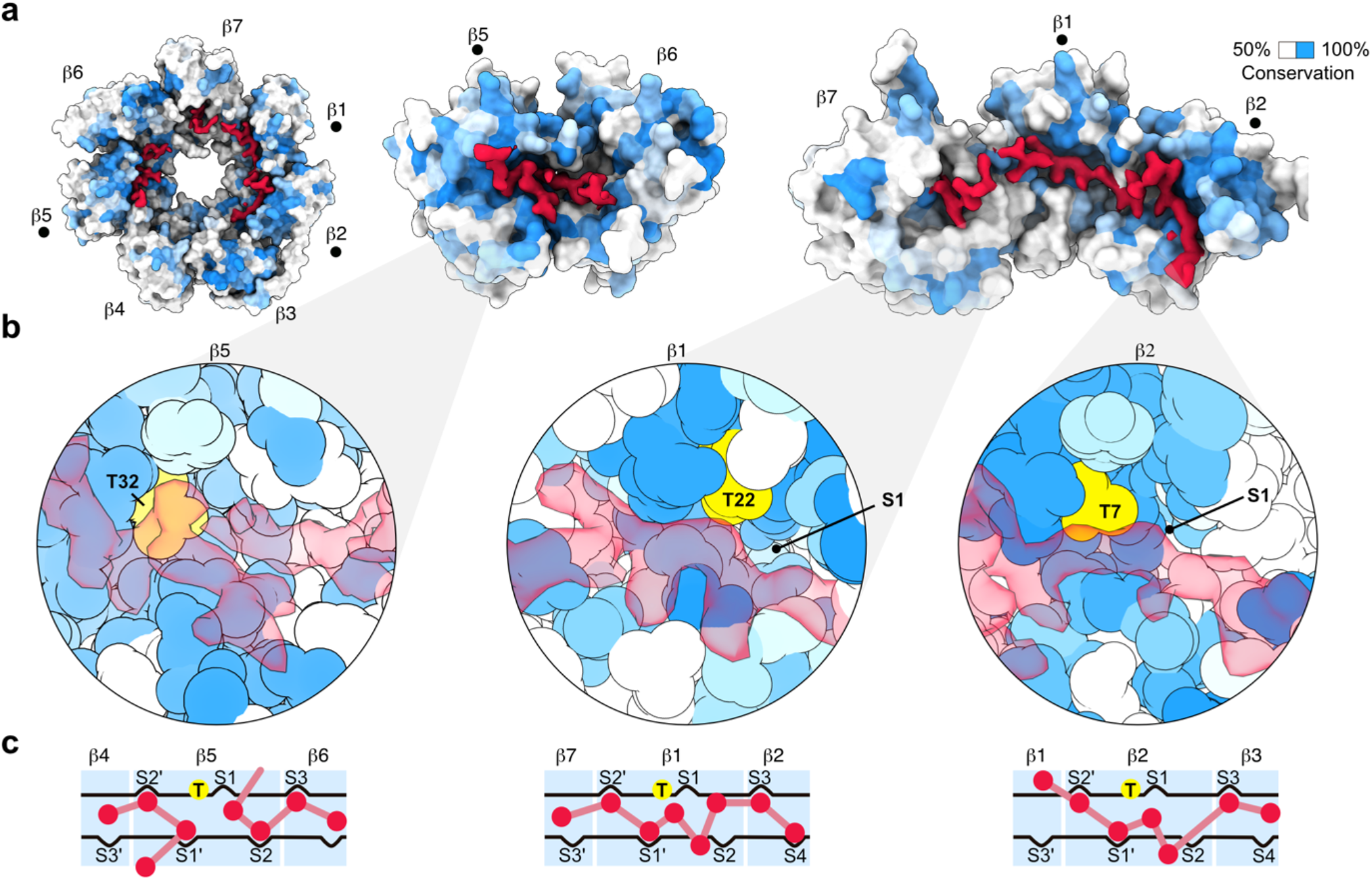
Spore proteasomes accumulate non-proteasomal peptide fragments in the proteolytic chamber. **a** Top-down view of one β-ring (left) next to selected peptide-bound β-subunits. The selected proteasome subunits are shown with surface representations, colored in shades of blue based on eukaryotic sequence conservation (from low conservation in white to high conservation in blue), superimposed with the cryo-EM densities of the non-proteasomal fragments (red). Subunits marked by a dot highlight those responsible for proteolysis. **b** Zoom-in sections of active sites of the proteolytically active β-subunits. The structure is shown as spheres colored according to conservation as in (a), with the active-site threonine highlighted in yellow and the non-proteasomal fragments as red transparent cryo-EM densities. **c** Schematic representation of proteolytic active sites and specificity sites, with active site threonines shown in yellow and density-derived peptide fragment orientations shown in red.

The cryo-EM map of the spore-isolated proteasome displays pronounced additional density in the central chamber (**Fig. 1c and 2a**). Multiple peptide-like densities are present, with the most prominent and continuous density found in the catalytic chamber. Each antechamber contains two poorly-resolved fragments, while each β-ring houses at least three resolved peptides: two shorter fragments of approximately 7-10 residues and one more extended fragment of roughly 30 residues (**Fig. 2a**). The two short fragments are bound to the β5 active sites and contact the inactive β6 subunit (**Fig. 2**). The more extended peptide originates from the inactive β7 subunit and travels along conserved residues near the active sites of β1 and β2 (**Fig. 2b**). In all cases, peptides are bound to the substrate grooves but adopt suboptimal conformations for proteolysis (**Fig. 2c**). The longer peptide occludes the β1 and β2 active sites and interacts with several of the substrate binding cavities adjacent to the active site threonines. However, it does not directly contact the S1 and S2 sites and the backbone trace runs ∼ 4 - 5 Å away from the nucleophilic hydroxyl group, suggesting these suboptimal substrate conformations are incompatible with proteolysis. Two shorter peptides are bound to either side of the active site threonine of β5, limiting interactions with other potential substrates. Previous work indicates that peptidic ligands bound to the active site of the β5 subunit induce minor but distinct conformational changes in β5 tertiary structure^31^. Our spore-derived β5 subunit is essentially identical to the sporoplasm-derived β5 subunit, which lacks additional peptidic density, indicating these peptide fragments do not induce conformational changes in β5. Interestingly, the equivalent peptide fragments in the two β-rings are nearly indistinguishable in volumes refined with and without C2 symmetry (**Supplementary Fig. 1**). This suggests that they correspond to either an indigestible peptide sequence from an abundant substrate degraded at the late intracellular sporogenesis phase, when nutrients become limiting, or to highly expressed short spore peptides with an affinity for the interior of the proteasome.

### Conservation and variation in 20S proteasome active sites

Proteasomes are N-terminal nucleophilic hydrolases that use a deprotonated threonine hydroxyl group to cleave peptide bonds. Activation of the Thr residues in subunits β1, β2, and β5 requires the cleavage of propeptides after highly conserved glycine residues^32^. Substrate preference varies between the three protease subunits and is determined by residues in each S1 specificity pocket^23^. Catalytic residues are identical in microsporidia and other eukaryotes, indicating that microsporidian proteasomes utilize a cleavage strategy similar to other eukaryotes. Additionally, the S1 specificity pockets are largely conserved, demonstrating sequence specificities of *V. necatrix* proteasomes mirror those of other eukaryotes.

Residues in adjacent β subunits contribute to the structure of S1 pockets and help form S3/S4 specificity sites, which are often targeted during the development of selective proteasome inhibitors^23,33^. This region is more divergent in *V. necatrix* (**Fig. 3**), with large changes to the charge distribution around the β2 (trypsin) site. The negatively charged S3/S4 specificity site residues, which stabilize the positively charged substrates, are largely replaced with neutral or positive residues, providing a potential target for the development of selective therapeutics. Notably, Met45 in the β5 S1 site is retracted, potentially due to van der Waals interactions with Tyr53 and His118 of β6^34^, creating a larger binding pocket (**Supplementary Fig. 3**). Previous work has established that the relative orientation of Met45 is important for the binding of several inhibitors^34,35^. In fact, several selective proteasome inhibitors like PR-957 have a 15-to 255-fold higher affinity for mammalian immunoproteasomes than for constitutive proteasomes^34,35^. The Met45 of immunoproteasomes is in an ‘open’ conformation, allowing the efficient insertion of a bulky sidechain into the S1 site. In constitutive proteasomes, on the other hand, Met45 is in a ‘closed’ conformation that sterically inhibits PR-957 interactions^34^. The retracted Met45 structure visible in *V. necatrix* proteasomes is consistent with this open conformation (**Supplementary Fig. 3**). PR-957 and related compounds may therefore represent a reservoir of known inhibitors useful for the selective targeting of microsporidian proteasomes. In contrast, several other classes of known inhibitors are unlikely to bind in microsporidia. Thiol-reactive maleimides, for example, were designed to covalently bind to a well-conserved Cys118 in β3^33,36^. The mutation of this residue to an alanine in *V. necatrix* may therefore result in a low affinity for this class of inhibitors.

**Figure 3.**
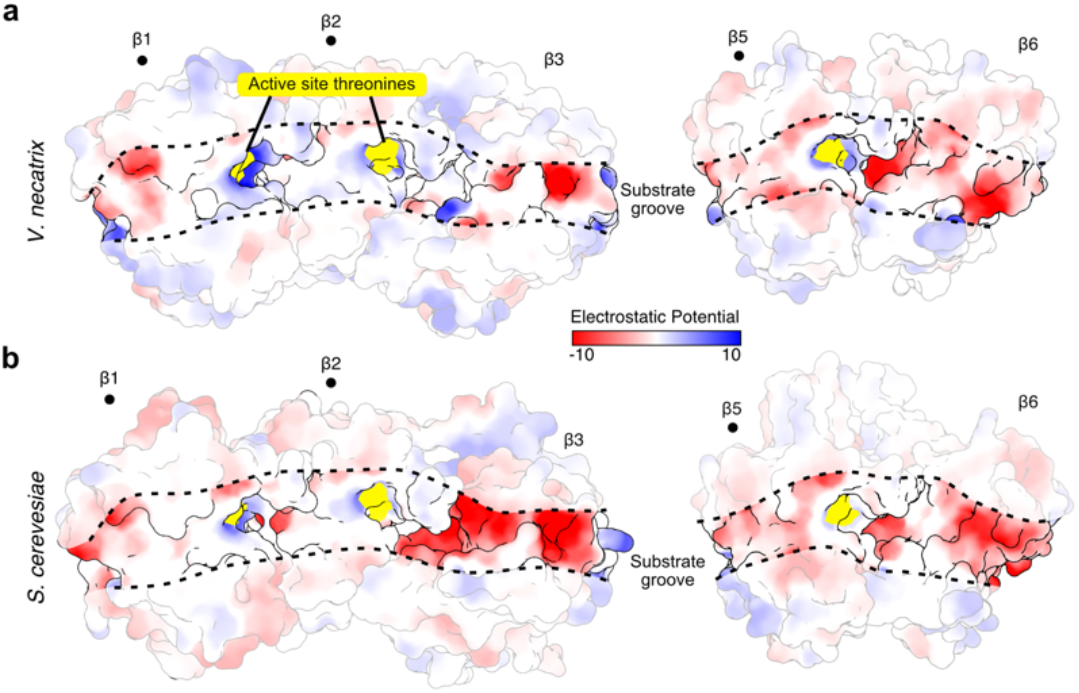
Electrostatic potential for substrate binding grooves. Surface representation of subunits important for substrate specificity in the proteolytic active sites of *V. necatrix* (**a**) or *S. cerevisiae* (**b**), colored based Coulombic electrostatic potential in kcal/(mol·e). Active site residues are colored in gold. Subunits harboring active sites are denoted with a •.

### Microsporidia evolved a different gate access mechanism

The substrate access to the proteolytic chamber is tightly controlled through gated pores in the α-rings^37^. Association of accessory proteins harboring C-terminal hydrophobic-tyrosine-X (HbYX) motifs, such as the RP, induces a conformational change in the N-terminal tails in the α-ring subunits^38^. A conserved YDR motif (**Fig. 4a**) is involved in the formation of the open-gate conformation through specific interactions between the tyrosine and a proline, present in all subunits^37^.

**Figure 4.**
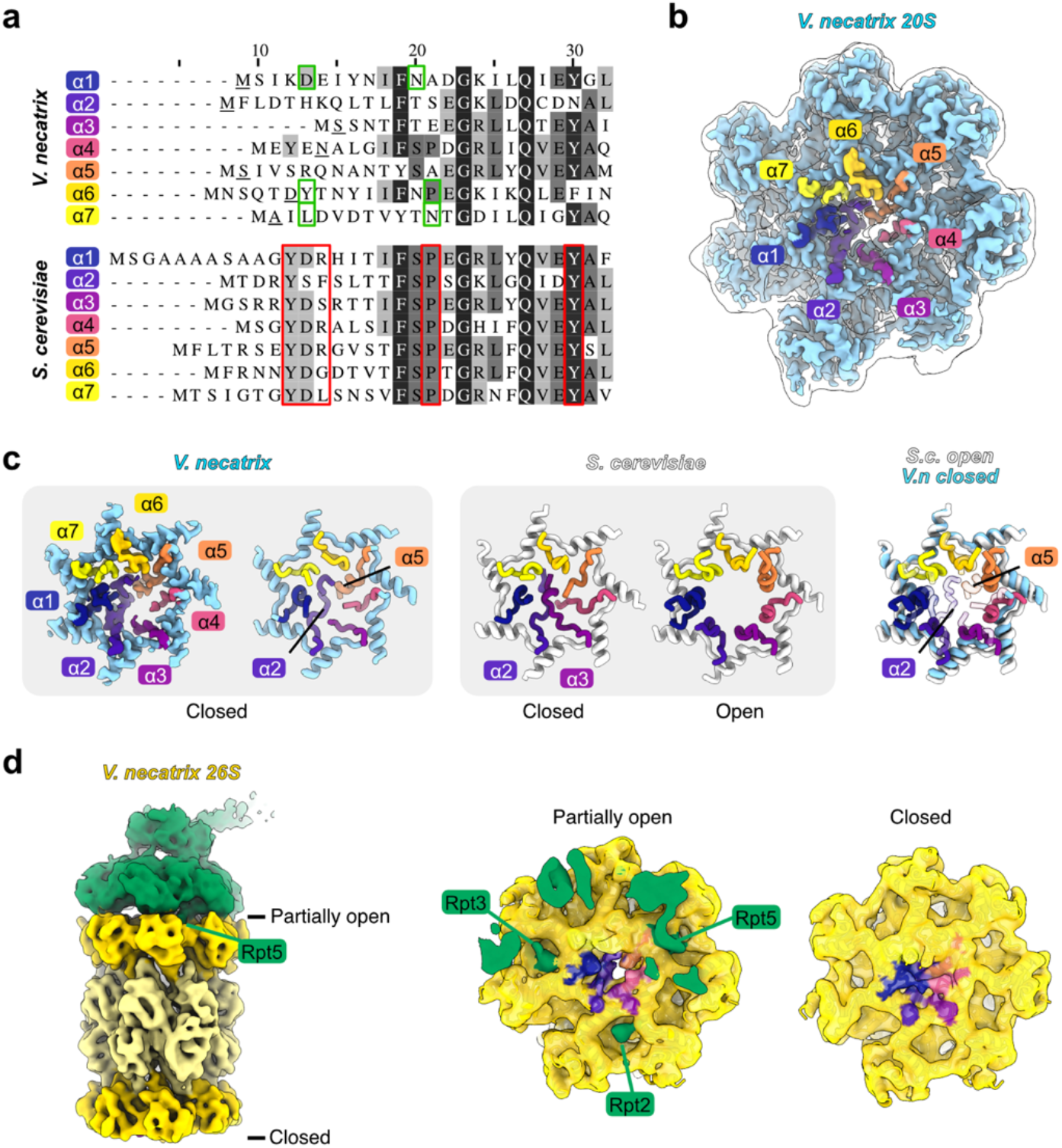
Microsporidia lack the conserved YDR motif and differ in gate lining *α*-subunit tail structure. **a** Alignment of the *V. necatrix* (top) and *S. cerevisiae* (bottom) *α*-ring subunit tail amino acid sequences. Critical residues for the open-gate formation in eukaryotes are lined with red borders, the first resolved residues in the *V. necatrix* cryo-EM density are underlined in black, and the residues involved in an open-gate like conformation in the 20S structure are boxed in green. Sequence conservation is indicated with shaded residues from white for low conservation to black for highly conserved positions. **b** The 20S cryo-EM map with the N-terminal tails, colored and labeled as in (a), is depicted in solid and superimposed with a transparent white lowpass-filtered map. **c** The cryo-EM density of the N-terminal tails next to the *V. necatrix* structural model and corresponding regions from *S. cerevisiae* (closed; PDB-5CZ4^32^, open; PDB-6EF3^75^) are shown in isolation. The right side depicts a superposition of the *S. cerevisiae* open-gate and the *V. necatrix* closed-gate pore region. **d** The ATPase module (green) and the core (shades of yellow) of the 26S cryo-EM density are shown in isolation next to a slice through the RP-CP interaction interface (middle) and a view from the bottom. The tail regions are colored as in (a-c).

Microsporidia have significantly changed the N-terminal tails of the α subunits (**Fig. 4a**) and evolved an alternative mechanism to restrict access. Overall, the tails are shorter than in yeast orthologs and adopt a different closed-gate conformation. In yeast, the N-terminal tails of multiple subunits (α2 to α5) extend into the pore and block the access. The tails of α2 and α4 descend into the pore while α3 and α5 sit on top (**Fig. 4b**). In *V. necatrix*, the closed gate is formed predominantly by the α2 and α5 tails. The tail of α2 takes over the function and space of yeast α2 and α3 tails, and a short helical segment in α5 serves as a plug. The pore of this gate is less rigorously closed, resulting in a small hole visible between chain α2 and α4 (**Fig. 4**). Further, the conserved YDR motif and the following proline, required to stabilize the open-gate conformation, have been lost in all chains except α6 (**Fig. 4a**). Although the characterized spore and sporoplasm 20S proteasome are in a closed-gate state, the N-terminal tails of α1, α6 and α7 can be compared to the yeast open-gate conformation (**Fig. 4b, c**). Despite the absence of RP, these tails adopt a conformation similar to the yeast open-gate proteasome. The terminal five residues of the tail of α6 are not well resolved, but the ensuing Tyr7 interacts with Pro15 and resembles the yeast open conformation. Similarly, despite lacking the tyrosine or the proline, α1 and α7 adopt open-gate-like conformations (**Fig. 4c**). For opening the gate, only two tails require remodeling, as opposed to four in yeast.

The C-terminal HbYX motifs of the ATPase subunits Rpt2, Rpt3, and Rpt5 are only partially retained in microsporidia (**Fig. 4d**), suggesting analogous, but non-identical, binding interactions between the RP and CP are present. Recent work has demonstrated the importance of other Rpt C-termini in the binding and gate-opening of the 26S proteasome^39^. While N- and C-terminal deletions are present in most microsporidian proteasomal orthologs, the Rpt subunits terminate at the same residue as their yeast orthologs, with the exception of a two-residue truncation in Rpt3 (**Fig. 4d**). These data support a conserved role for the C-termini of Rpt proteins in binding and gate-opening of microsporidian proteasomes.

### The architecture of the microsporidian 26S proteasome

Proteasomes isolated from germinated sporoplasms were capped by RPs in about 12% of all particles, and a small fraction was capped on both ends. Despite limitations in starting material and the small total population of capped proteasomes, we were able to generate a low resolution (8.3 Å) map of the 26S proteasome from *V. necatrix*. To structurally characterize the complex, we identified microsporidian orthologs, predicted tertiary structures using AlphaFold^40^, and superimposed models onto the closest homologous structure from yeast. Models were then manually adjusted to best fit the volume. *Vairimorpha necatrix* proteasomes display a similar general structure to 26S proteasomes from other eukaryotes (**Fig. 5a**), with the barrel-like CP capped by a ring of six ATPases and a “grasping-hand”-like lid complex^41^. Interestingly, there is a conspicuous lack of density for three RP subunits: Rpn12, Rpn13, and Rpn15 (**Fig. 5b**). Although Rpn13 is often poorly-resolved^41^, our findings are corroborated by previous work that was unable to identify microsporidian orthologs for these proteins^27^. In contrast, clear density for Rpn3 is visible, and a highly truncated version of Rpn3 is identifiable using BLAST ^42^ (**Fig. 6, Supplementary Data 2**), despite prior speculation that it is absent in microsporidia^27^.

**Figure 5.**
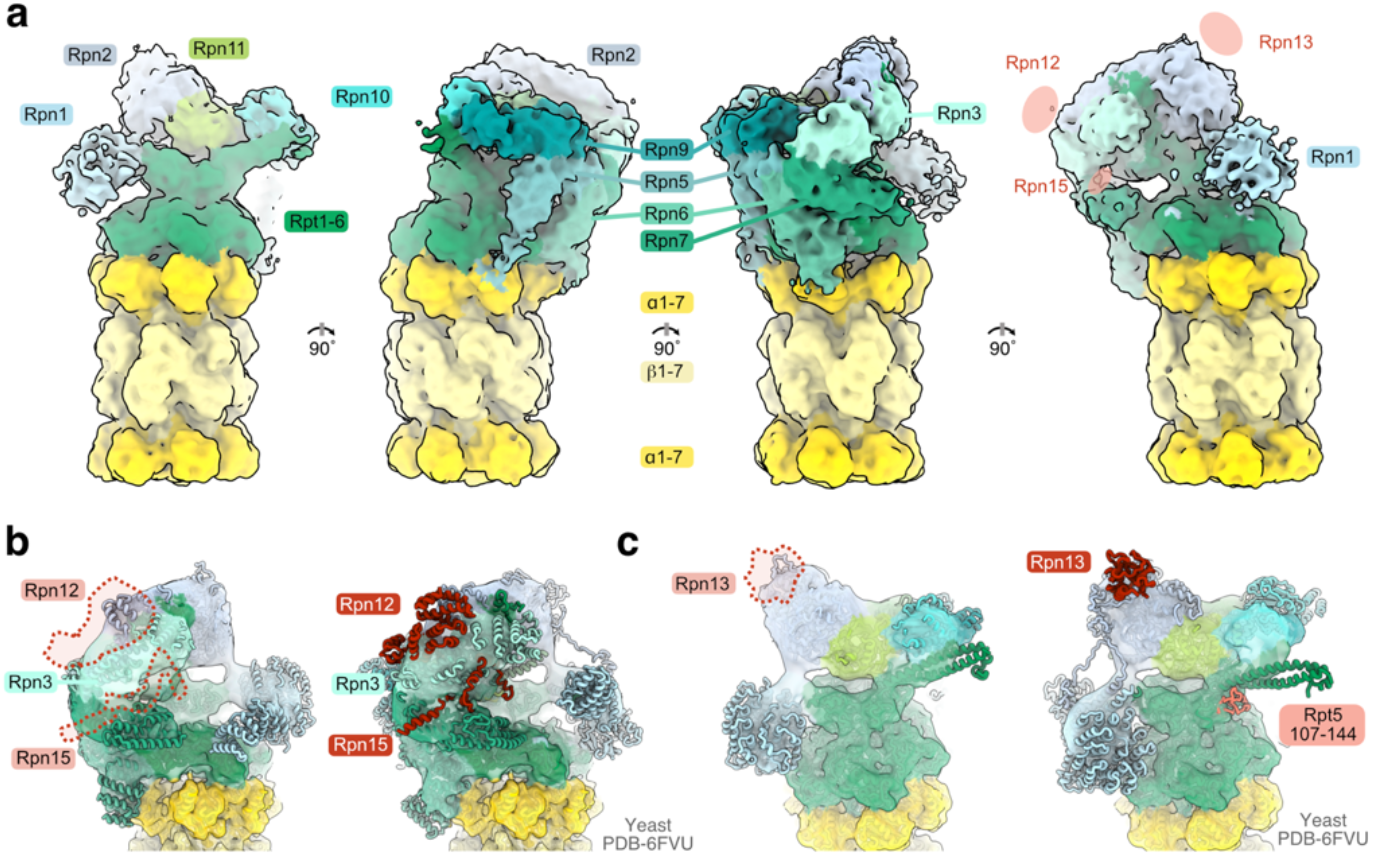
The architecture of the microsporidian 26S proteasome and missing regulatory subunits. **a** Four 90°-rotations of the sporoplasm-isolated 26S cryo-EM density are shown with the regulatory particle components colored in shades of green and the core particle indicated in shades of yellow. The location of eukaryotic subunits not observed in the microsporidian proteasome density is indicated in the right-most view. **b**,**c** Two different views of the microsporidian regulatory particle cryo-EM density are shown with docked AlphaFold^40^ models (left) and the *S. cerevisiae* structure (PDB-6VFU^39^). Missing elements are indicated with dashed lines and colored red in the yeast structure.

**Figure 6.**
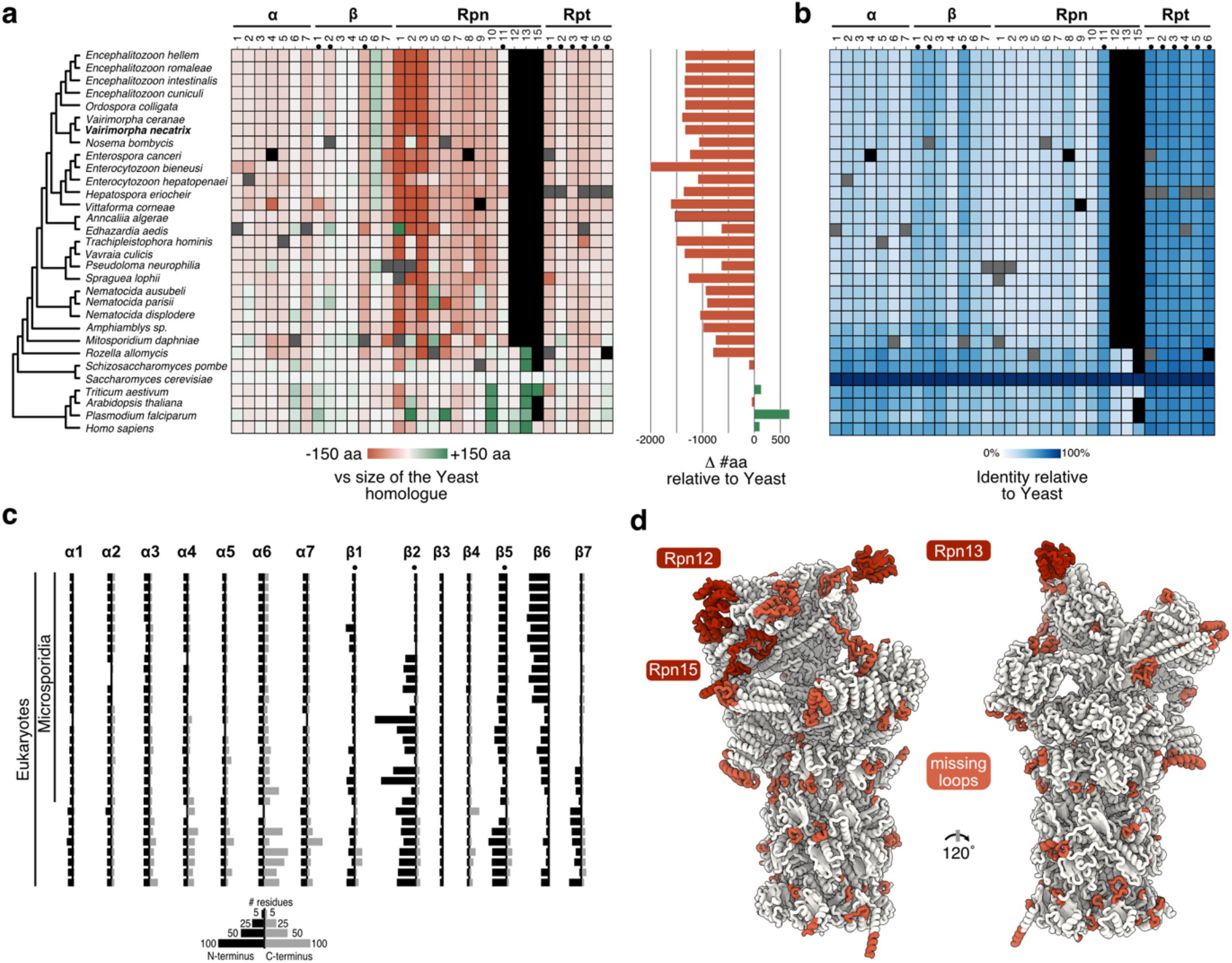
Phylogeny and reductive evolution of the microsporidian proteasome. **a** Phylogenetic tree of proteasome core and regulatory subunit sequences of microsporidia and outgroups. The plot shows the sequence length of each protein (left) as well as the total proteasome sequence length (right) compared to *S. cerevisiae* (red: shorter, green: longer, grey: incomplete sequence, black: not clearly identifiable). **b** Heat map of the protein sequence identity of each subunit compared to *S. cerevisiae* (white: lowest sequence identity, dark blue: highest sequence identity, grey: incomplete sequence, black: not clearly identifiable). **c** Histogram plots of the number of N-(grey bars) and C-terminal residue (black bars) from a defined position in the multiple sequence alignment of the individual subunits. **a-c** A black dot on top marks enzymatically active subunits. **d** Two views of the *S. cerevisiae* 26S structure (PDB-6FVU^39^) colored in white with the subunits Rpn12, Rpn13, and Rpn15, which are absent in microsporidia, and missing internal and terminal elements colored in shades of red.

The truncation of Rpn3 and absence of Rpn12/15 alters the overall lid structure, and the “grasping hand” in microsporidian proteasomes has a smaller thumb and a much more open purlicue (**Fig. 5**). It is unclear how this more open structure affects enzymatic activity or proteasome stability; however, there is a surprisingly minor effect on proteasome structure in other regions. For example, in most eukaryotes the C-terminal tails of the eight structural lid proteins (Rpn3/5/6/7/8/9/11/12) all congregate in a central and essential helical bundle^43^. The final helix integrated, that of Rpn12, induces widespread structural remodeling of the lid and elicits the formation of the complete RP complex^44^. The absence of Rpn12 in *V. necatrix* precludes the integration of its helix. Nonetheless, the proteasome adopts a conformational state akin to the final remodeled structure in yeast and displays clear central density that likely represents the helical bundle.

Reductive evolution has also affected subunits responsible for the recognition and deubiquitination of substrates. Rpn13, one of three ubiquitin receptors in the RP, is absent in our structure and is not identifiable in any microsporidian species (**Fig. 5, Supplementary Data 2**)^27^. Deletions of Rpn13 in yeast are associated with a 50% decrease in binding to ubiquitin-conjugated substrates^45^, suggesting this deletion reduces the ubiquitin binding potential of microsporidian proteasomes. Additionally, the well-conserved ‘N-loop’ of Rpt5 has been removed (**Fig. 5c**). This loop directly interacts with ubiquitin-bound Rpn11, stabilizing the interaction and optimizing the positioning of the isopeptide bond for efficient deubiquitylation^46^. These examples demonstrate that microsporidia have streamlined proteasome structure, and removed segments associated with important, but nonessential functions.

A comparison of the quaternary arrangement of *V. necatrix* subunits to six known yeast conformational states indicates our structure is most consistent with the s2 state^39^. Proteasomes in the s2 state are considered primed for degradation, as the CP, ATPase ring, and Rpn11 are appropriately aligned for substrate deubiquitinating and eventual degradation. The CP gate is largely closed in the yeast s2 conformation, leaving a gap of only ∼5 Å that hinders access to the catalytic chamber^39,47^. The opened s4 gate, in comparison, is approximately 17 Å wide. Although the majority of our structure is reminiscent of the yeast s2 state, the gate occupies an intermediate conformation between opened and closed, with a gap diameter of approximately 11 Å. The C-terminal HbYX motifs of Rpt2, Rpt3, and Rpt5 are docked within the α-ring of the CP (**Fig. 4d**), as is typical^38^, despite the small C-terminal truncation in Rpt3 and mutations in the Rpt5 motif. The C-termini of Rpt1 and Rpt6, known to induce gate opening^39^, are not visibly buried within the α-ring. These data together point to the isolated *V. necatrix* proteasome adopting a primed, but inactive conformation. The incomplete obstruction by the CP gate may allow for a more passive degradation of substrates, allowing microsporidia to conserve vital ATP.

### Phylogeny and reductive evolution of microsporidian proteasomes

The taxonomic placement and organization of microsporidia has undergone significant revisions due to the absence or reduction of several organelles and the remarkable sequence variation between species. Indeed, two evolutionarily adjacent *Nematocida* spp. share only ∼68% of their sequence^48^, and even in complexes as universally conserved as ribosomes, yeast and microsporidian homologs are only 38% identical in sequence on average^3^. This sequence diversity confounds not only phylogenetic classification, but also attempts to identify microsporidian homologs, which hampers our functional understanding of these ecologically important organisms. Several clear examples can be seen in previous work, where ribosomal and proteasomal proteins were unidentifiable using BLAST^27,49^. Structural studies revealed that several of these proteins are retained^1,12,14^, albeit with low sequence conservation. In our own work, initial attempts to identify *V. necatrix* α subunits resulted in nearly identical top hits for α1, α4, and α5. Only by modelling the chains into density were we able to ascertain the correct sequences, reinforcing the importance of structural data as a complementary tool to assist in the annotation and functional characterization of microsporidian complexes.

We have identified microsporidian proteasome subunits on MicrosporidiaDB^50^ and NCBI^42^ using a combination of yeast sequences and structurally validated *V. necatrix* proteins as queries (**Supplementary Data 2**). The phylogenetic tree generated using high-confidence proteasomal subunits (**Fig. 6a**) is consistent with trees derived from simplified rRNA or whole-proteome analyses ^3,51^. Saliently, a comparison of the relative length and conservation of proteasomal subunits between microsporidia and selected eukaryotes to *S. cerevisiae* reveals widespread evidence of reductive evolution (**Fig. 6a**).

Although the functions of many eliminated fragments are not well-defined, some notable examples include the poorly resolved^52^ C-terminal extensions from α3, α4, α5, and α7, which are associated with stabilization of CP-RP interactions, substrate recognition^53^, nuclear localization^52^, and recruitment of the quality control protein Ecm29^54^. The propeptides of the β subunits are also frequently modified. These segments are integral to the assembly process, where they protect catalytic residues and recruit other subunits and assembly factors^55,56^. The most consistent and sizable truncations occur within β5 and β7 propeptides (**Fig. 6c**), which recruit β6 and promote proteasome dimerization via interactions with Ump1, respectively^55,56^. Contrary to the tendency to extreme reduction, many microsporidians have expanded their β6 propeptides by more than 70 amino acids. These substantial changes to microsporidian propeptides suggest that the proteasome assembly pathway differs significantly from typical eukaryotes. Thus, while microsporidia express asymmetric proteasomes that are evolutionarily related to eukaryotes, the lengths and structural variability of α and β subunits are converging towards prokaryotic parameters. The average sequence lengths of *V. necatrix* α (233 ± 5) and mature β (204 ± 10) subunits are very similar to the lengths of α (233) and β (203) chains from the model archaeon *Thermoplasma acidophilum*. This is significantly smaller than the size and variability present in yeast α (257 ± 16) and mature β (214 ± 15) chains.

Microsporidia seem to have universally lost Rpn12, Rpn13, and Rpn15, and almost all remaining proteins have been pared down, resulting in proteasomes approximately 1500 amino acids smaller than in yeast. Additionally, structural subunits display a marked decrease in sequence conservation compared to enzymatic subunits (**Fig. 6b**). Large deletions are particularly common in the RP, with the Rpn1, Rpn2, and Rpn3 deletions exceeding 100 amino acids each.

In some cases, the significance of a deletion is readily apparent. For example, truncation of the ∼80 C-terminal amino acids of Rpn2 coincides with the absence of Rpn13, known to bind to this region^57^. This observation is consistent with a previous hypothesis that the elimination of proteins in microsporidia has resulted in a simplified interaction network, which in turn enables the erasure of protein-interacting domains^58^. Likewise, the N-terminal ∼100 amino acids of Rpn3 protrude from the proteasome lid in yeast and may stabilize the Rpn12 binding region. The loss of Rpn12 is thus accompanied by the loss of these Rpn3 residues in microsporidia. In Rpn1, deletions may denote regions where binding partners have yet to be identified. The approximately 100 amino acid deletion in Rpn1 connects two adjacent toroid domains in other eukaryotes but is absent from all reconstructions due to its structural heterogeneity^59^. Widespread conservation of this disordered segment in other eukaryotes signifies an important functional role, such as the potential recruitment of an unknown partner, which may be absent in microsporidia.

Although it is tempting to draw conclusions from the absence or extreme modification of several microsporidian orthologs, isolated outliers are most likely the result of incomplete genome assemblies and further work is necessary to establish the veracity of any remarkable modifications. On the other hand, alterations present in a cluster of organisms, like the expansion of Rpn6 in *M. daphinae* and *Nematocida* spp. (**Fig. 6a**), may be worth investigating to identify the potentially expanded repertoire of proteasome functionalities.

## Discussion

Stringent reductive evolution in microsporidia has eliminated many genes required by free-living species, including those necessary for sugar/fat metabolism, amino acid/nucleotide biosynthesis, and energy generation via oxidative phosphorylation^60^. Microsporidia are thus left with a core of only 800 conserved proteins, in addition to species and clade specific proteins^16^. Proteins that are retained are involved in vital cellular processes like protein synthesis and recycling^58^. The extreme reduction of microsporidian proteins and protein complexes makes microsporidia ideal organisms in which to study minimized versions of essential and highly conserved macromolecular assemblies. Previous work on the ribosome has demonstrated that microsporidia have rolled back many of the expansions characteristic of eukaryotic ribosomes and deleted several proteins known to be dispensable in yeast^1,12,14^. These data suggest that microsporidian complexes can be used to highlight regions with nonessential or expanded functionalities in eukaryotes.

The 20S structure described in this work is the most minimized eukaryotic proteasome studied to date and reveals that microsporidia have deleted many of the insertions that differentiate individual α and β subunits. Overall, the 26S proteasome is approximately 170 kDa smaller in *V. necatrix* than in yeast (**Fig. 6**), with nearly every subunit undergoing some level of reduction. Truncations most frequently occur in disordered loops or at the N- and C-termini. As short motifs used for the recognition of interaction partners often occur within disordered segments, the deletion of these regions results in a much more simplified proteasome interaction network when compared to other eukaryotes. For example, the deletion of the ubiquitin receptor Rpn13 coincides with the removal of complimentary interacting interfaces on Rpn2 and the deletion of binding partners like Uch37, which binds to the C-terminal region of Rpn13^61^. Uch37 is a deubiquitinase that serves a proofreading function by clearing improperly or inadvertently ubiquitylated substrates. Interestingly, previous studies on translation demonstrated that microsporidia have also removed several proofreading functionalities from ribosomes^12,62^, suggesting that such editing or high-fidelity tasks are common targets for macromolecular streamlining in microsporidia.

Unlike bacterial and archaeal proteasomes, which can self-assemble, eukaryotic proteasomes require a diverse group of assembly factors to assist in the formation of mature particles^63^. Core particle assembly in eukaryotes is largely mediated by Pba1-4, Blm10, Ump1, and the propeptides of the β subunits. As in a previous study^27^, we were unable to identify any of these assembly factors in most microsporidians. This, in combination with the frequent changes to the propeptides, suggests that microsporidia utilize a novel assembly pathway or are able to self-assemble. Intriguingly, the β6 subunit is the only proteasomal protein expanded in most microsporidian species, due to extension of the propeptide (**Fig. 6c**). One possibility is that β6 has cannibalized the roles of the truncated subunits by incorporating assembly factor binding sites into the β6 propeptide, and the low sequence identity inherent to microsporidia has made identification of assembly factors difficult. Alternatively, microsporidian proteasomes may be amenable to self-assembly, like bacterial and archaeal equivalents, with β6 playing a larger role in the assembly process. These changes facilitate simplification of the proteasome interaction network, allowing microsporidia to conserve resources by eliminating unnecessary genes.

Proteasomes account for 5% of the total protein content in yeast^64^. They therefore represent a large investment of nutrients and require consistent ATP availability to function. Microsporidian spores display very little background metabolic activity and are still infective after a year in environmental conditions^65^. It is thus imperative for microsporidia to conserve energy and nutrients during the extracellular spore stage, necessitating a means to impede proteasome activity. Previous studies have confirmed that microsporidian spores are rich in proteasomal proteins, indicating they are not digested during the extracellular stage^28^; however, it was unknown how microsporidia regulate proteasomal activity. To compare potential macromolecular assemblies from ATP-poor extracellular spores and ATP-rich intracellular sporoplasms, we purified proteasomes from dormant and germinated spores in the absence or presence of ATP, respectively. Proteasomes isolated in ATP-rich conditions formed full 26S assemblies. On the other hand, proteasomes from dormant spores, isolated without ATP, formed only 20S core particles and displayed additional peptide density within the CPs (**Fig. 1**).

Although we were unable to unambiguously identify the peptide fragments bound to the active sites of the 20S proteasome, the orientation of their binding appears to be inconsistent with active digestion (**Fig. 2**). One possibility is that these fragments are derived from proteins highly abundant in either the spore or the sporont stage and are the final proteins digested before quiescence, leading to high occupancy in the 20S proteasomes. Alternatively, the additional density may represent previously undescribed dormancy factors, as has been noted for other microsporidian macromolecular assemblies^1,12^. As dormancy factors, these fragments could sterically inhibit low-level background digestion by proteasomes during the spore stage. We speculate that, upon exposure to ATP, binding of the RP to the CP leads to the dissociation of the bound peptides, enabling proteolytic digestion. This mechanism would allow for the rapid and efficient reactivation of proteasomes post-infection of host cells, while facilitating the conservation of nutrients during the spore stage. Interestingly, this process may be reminiscent of the PSG-based storage pathways present in other eukaryotes during carbon starvation, where nutrients are conserved by sequestering proteasomes rather than degrading them^19,21^.

Inhibition of proteasomes can lead to the accumulation of misfolded proteins and cell death. Prior studies in microsporidia have suggested that they are particularly sensitive to proteotoxic stress due to limitations imposed by genome compaction^27^. Although many structural features of the proteasome are altered in microsporidia, the essential enzymatic folds are conserved (**Fig. 6b**). Both the active site residues and the physicochemical properties of the substrate binding pockets are largely maintained, suggesting that many known inhibitors that target proteolytic sites will function in microsporidia. Interestingly, the β5 active site adopts a conformation associated with selective binding by a class of peptidomimetic inhibitors^35^, reinforcing the feasibility of proteasome inhibitors as potential therapeutics against these emerging pathogens. Additionally, the genomic streamlining in microsporidia has resulted in a highly differentiated and minimal version of eukaryotic proteasomes, revealing unique interaction interfaces that may be utilized in future studies to serve as drug targets.

## Methods

### Cultivation and isolation of *V. necatrix*

*V. necatrix* was cultivated and reproduced by feeding approximately 100,000 spores to fourth and fifth instar *Helicoverpa zea* larvae, grown on a defined diet (Benzon Research). After three weeks at 21–25 °C, the spores were harvested. First, larvae were homogenized in water, followed by filtration through two layers of cheesecloth and subsequent filtration through a 50 μm nylon mesh. The filtrate was layered on top of a 50% Percoll cushion in a 2-ml microcentrifuge tube, and spores were pelleted via centrifugation at 1,000*g* for 10 minutes. The pure spores were stored at - 80 °C until further use.

### Purification of the *V. necatrix* proteasome

Proteasomes were purified from either dormant or germinated spores to identify differences between inactive and activated complexes. For spore-based preparations, 100 mg of *V. necatrix* spores were suspended in 500 μl size exclusion chromatography (SEC) buffer containing 50 mM Tris (pH 7.4) and 300 mM NaCl, supplemented with 5 mM DTT and a protease inhibitor cocktail (Complete EDTA-free, Roche). Spores were transferred to tubes containing Lysing Matrix E (MP Bio), and lysed via three, one-minute intervals in a Fast-Prep 24 (MP Bio) grinder at 5.5 m/s. Complete lysis was guaranteed by a following sonication step and verified via light microscopy. Spore debris was then pelleted via centrifugation for 20 minutes at 20,000*g*, before loading the supernatant onto a Superose 6 Increase 10/300 SEC column (Cytiva). Fractions containing proteasomes were identified using a combination of SDS-PAGE and negative stain EM.

To obtain sporoplasm-derived proteasomes, 5 mg of *V. necatrix* spores were germinated via alkaline priming^29^. Briefly, spores were incubated in 500 μl of 0.1 M KOH for 15 minutes at 22°C, followed by pelleting via centrifugation at 10,000*g* for 2 minutes. Primed spores were then germinated by resuspension in 650 μl of 0.17 mM KCl, 1 mM Tris-HCl (pH 8), and 10 mM EDTA. Germination was confirmed by light microscopy. Approximately 80% of spores germinated within 5 minutes; following which, 650 μl of rich insect cell media (ExCell 420) supplemented with 20 mM ATP, was added to the germination mix. Sporoplasms were incubated for 20 hours at 22°C, then pelleted via centrifugation at 10,000*g* for 2 minutes and resuspended in 500 μl proteasome buffer containing 50 mM Tris (pH 7.4), 100 mM NaCl, 10m M MgCl2, 20 mM ATP, 1m M DTT, and 1% Glycerol. Sporoplasms were then lysed via mild sonication (6 μm amplitude) for a total for 12 seconds, and lysis was validated using light microscopy. Purification of proteasomes proceeded via differential centrifugation in proteasome buffer, with sequential hour-long spins at 21,000*g*, 54,000*g*, and 121,000*g* at 4°C. Pellets were then resuspended in 50 μl proteasome buffer without glycerol. Negative stain EM and SDS-PAGE were used to verify 20S and 26S proteasome enrichment in the 121,000*g* fraction.

### Phylogenetic analysis

The proteasome sequences were retrieved by tblastn searches from NCBI^42^, MicrosporidiaDB^50^, and an in-house *V. necatrix* genome (manuscript in preparation), using *S. cerevisiae* and *V. necatrix* sequences as queries with an E-value cutoff of 0.05. The phylogenetic tree was generated by protein sequence alignment using MUSCLE5 (v5.1.)^66^. Alignments were trimmed using trimAl (v1.4.1.)^67^ with the – gappyout option. FASconCAT-G (v1.05.1.)^68^ was used to concatenate the trimmed alignments and to create the partition file. The phylogenetic tree was constructed using iqtree (v2.2.0.)^69^ with the -MFP MERGE function to identify the best partition model, followed by tree reconstruction using 1000 bootstrap replicates. The -p function was used to allow each partition to have its own evolution rate. Tree rooting was done in Figtree (http://tree.bio.ed.ac.uk/software/figtree/, v.1.4.4.). The sequence identity heatmap was constructed using MUSCLE5 (v5.1.) ^66^ with *S. cerevisiae* sequences set as reference.

### Cryo-EM grid preparation and data collection

Purified proteasomes were applied as 3.5 μl aliquots to Quantifoil R2/1 200-mesh copper grids (EM sciences, Prod. No. Q250CR1) or R1.2/1.3 400-mesh gold grids with 2-nm carbon (EM sciences, Prod. No. Q450AR1.3-2nm), for spore and sporoplasm samples, respectively. Grids were glow discharged for 30 seconds at 15 mA before sample application. Grids were blotted for 5 seconds in an FEI Vitrobot Mark IV (Thermo Fisher Scientific), set to 4°C and 100% humidity, prior to plunge-freezing into liquid ethane.

Cryo-EM data were collected at the Umeå Core Facility for Electron Microscopy. For 20S structures, data were collected with a pixel size of 1.042 Å on a Titan Krios (Thermo Fisher Scientific) operated at 300 kV using a Gatan K2 BioQuantum direct electron detector. Data for the 26S structure was collected at a 1.495 Å pixel size, using a 200 kV Glacios system (Thermo Fisher Scientific) equipped with a Falcon 4i direct electron detector. A total of four EPU data collections were used to generate the structures for spore-derived 20S, sporoplasm-derived 20S, and sporoplasm-derived 26S structures. Data collection statistics are summarized in **Supplementary Table 1**.

### Cryo-EM image processing

The four datasets were processed using a combination of Relion-3.1^70^, Relion-4.0^71^, and cryoSPARC v3.3.2^72^. The procedure is outlined in **Supplementary Fig. 1**. Briefly, in all datasets, movie alignments, drift correction, and dose weighting were done using MotionCor2^73^, and CTF estimation was performed with CTFFIND-4.1.14^74^. Micrographs with poor CTF fits or non-ideal ice thickness were removed, resulting in 3,935 micrographs for the spore-derived dataset one, 3,416 micrographs for the spore-derived dataset two, 3,277 micrographs for the sporoplasm-derived 20S data set, and 2,658 micrographs for the 26S dataset (**Supplementary Table 1**). For the two spore datasets, particles were first autopicked in Relion-3.1 using a Gaussian-blob template. Particles were classified in 2D, and promising classes were then used as templates for a second round of automated particle picking. Template-picked particles were extracted with a box size of 400 pixels (416 Å), filtered via iterative 2D and 3D classification, and further refined via per-particle CTF refinement and Bayesian Polishing in Relion-3.1. Polished particles were then exported to cryoSPARC and homogeneously refined, with C2 symmetry imposed. Map resolution was determined by the Fourier shell correlation (FSC) between two half-maps at a value of 0.143, resulting in a nominal 20S resolution of 2.77 Å (52,679 particles).

Micrographs from the sporoplasm-derived samples were manually picked in Relion-4.0. The 20S map was refined using the 1.042 Å pixel size dataset. Manually picked particles were extracted with a 400-pixel box size (416 Å) before exporting them to cryoSPARC for *ab initio* modelling and homogeneous refinement with imposed C2 symmetry, leading to a 3.2 Å map (17,942 particles). To offset the low 26S particle density, a 1.495 Å pixel size dataset was collected. Manually picked particles were again extracted with a box size of 400 pixels (598 Å), filtered via 3D classification, and non-uniformly refined without symmetry constraints. The final 26S map has a nominal resolution of 8.3 Å (6,442 particles) at an FSC value of 0.143.

### Model building and refinement

A high-resolution yeast proteasome crystal structure (PDB-5CZ4^32^) was used as an initial template for modelling *V. necatrix* proteasomes in Coot^75^. Model geometries and fits to maps were adjusted in ISOLDE^76^. Final refinements were performed using PHENIX^77^ (version 1.20-4487) real space refinement against the final map, resolved to 2.8 Å. Model statistics are described in **Supplementary Table 1**, and model compositions are described in **Supplementary Table 2**. Sidechains in poorly resolved areas were removed due to insufficient data. Structures and maps were visualized and presented using ChimeraX^78^.

## Supporting information

Description of Supplementary Data

## Data availability

The cryo-EM density maps and the coordinates for the microsporidian proteasome have been deposited in the EM Data Bank with accession code **EMD-15365**/**PDB-8ADN** for the spore stage 20S core-particle, **EMD-15367** for the sporoplasm 20S core-particle (post-germination), and **EMD-15366** for the full proteasome.

## Acknowledgments

We thank B. Vossbrinck for her help in editing the manuscript, SV Pipaliya for phylogenetic advice, and members of the Barandun laboratory for discussions and critical reading of this manuscript. Further, we thank Michael Hall and Camilla Holmlund for help with cryo-EM data collection. The electron microscopy data was collected at the Umeå Core Facility for Electron Microscopy, a node of the Cryo-EM Swedish National Facility, funded by the Knut and Alice Wallenberg, Family Erling Persson and Kempe Foundations, SciLifeLab, Stockholm University and Umeå University. N.J. is supported by an Integrated Structural Biology fellowship from Kempe (JCK-1918). J.B. acknowledges funding from the Swedish Research Council (2019-02011), the European Research Council (ERC Starting Grant PolTube 948655), the SciLifeLab National Fellows program, and MIMS. C.R.V. acknowledges funding from the Hatch Grant Project CONH00786 and R. Tyler Huning. The computations were enabled by resources provided by the Swedish National Infrastructure for Computing (SNIC) at High-Performance Computing Center North (Project Nr. SNIC 2021/23-718 and SNIC 2021/22-936), partially funded by the Swedish Research Council through grant agreement no. 2018-05973.

## Contributions

N.J. and J.B. conceived the study. C.R.V. cultivated microsporidia. N.J. purified proteasomes and performed all EM work. N.J. and K.E. determined the cryo-EM structures and together with R.R.W. built the structural model. R.R.W. and D.S. performed phylogenetic analysis and all authors interpreted the results, wrote, and edited the manuscript.

