## Supplementary Information for "Reductive evolution in the structure of the microsporidian proteasome"

**Supplementary Table 1.** Cryo-EM data collection, refinement, and model statistics.
**Supplementary Table 2.** *V. necatrix* proteasome subunit sequences and model composition.

**Supplementary Figure 1.** Cryo-EM data processing scheme.
**Supplementary Figure 2**. Overall and local resolution estimation.
**Supplementary Figure 3**. Conformational variation of the Met45 residue in β5 subunits.

**Supplementary Table 1. Cryo-EM data collection, refinement, and model statistics.**

| | **20S Spores** | **20S Sporoplasm** | **26S Sporoplas[illegible]** |
|---|---|---|---|
| | EMD-15365<br>PDB- 8ADN | EMD-15367 | EMD-15366 |
| **Data collection and processing** | | | |
| Voltage (kV) | 300 | 300 | 200 |
| Pixel Size (Å) | 1.042 | 1.042 | 1.495 |
| Electron exposure (e-/Å2) | 35.01 | 42.31 | 40.0 |
| Defocus range (um) | 0.7 – 2.8 µm | 1.2 - 3.0 µm | 1.2 - 3.0 µm |
| Symmetry imposed | C2 | C2 | C1 |
| Final particle images | 52,679 | 17,942 | 6,442 |
| Resolution (Å) | 2.8 | 3.2 | 8.3 |
| FSC threshold | 0.143 | 0.143 | 0.143 |
| Map sharpening B-Factor (Å2) | 68.3 | 66.1 | 514.1 |
| **Refinement** | | | |
| Initial model used | 5CZ4 | | |
| Model composition | | | |
| Non hydrogen Atoms | 47,368 | | |
| Protein residues | 6,041 | | |
| R.m.s deviations | | | |
| Bond length (Å) | 0.006 | | |
| Angles (°) | 0.961 | | |
| Validation | | | |
| MolProbity score | 1.48 | | |
| Clashscore | 5.75 | | |
| Poor rotamers (%) | 0.75 | | |
| Ramachandran | | | |
| Favored (%) | 97.06 | | |
| Allowed (%) | 2.94 | | |
| Outliers (%) | 0.00 | | |
| Ramachandran plot Z-scores | | | |
| whole | 0.69 | | |
| helix | 1.47 | | |
| sheet | 0.41 | | |
| loop | -0.36 | | |

**Supplementary Table 2.** ***V. necatrix*** **proteasome subunit sequences and model composition.**

| Subunit | Chains *SeqIDs* | Sequence (Propetide, Active site threonine, Sequence not observed, Modelled, Sidechains trimmed) |
|---|---|---|
| PSA1 | G,U<br>*H1G,H2U* | MSIKDEIYNIFNADGKILQIEYGLEAVNKSLPLVVLKNKNMIVCAAKKNQGHLLEDEVQTSFQPIYPNLYSA<br>FTGNWADVFYVNSKAKDLAHYASYKLGFSVTPDILCRKLADELQPLIQSTGERAPAFAGALFGFDNGKPVVY<br>MTNISAVCYPVYGSMVGSKNQNMYKYVEKYYNEDIEDEKLFEVAVGGLLESLGENSVYQEMEVAYLRNGEVL<br>KYLDDKEIESLLLSIADK |
| PSA2 | A,O<br>*H1A,H2O* | MFLDTHKQLTLFTSEGKLDQCDNALKAATQGSLSVGCVSENGVVLASLKESNNLVILSEYKKIYQISPNLGI<br>TYSGCQPDFRIQYNLSLKISEEYTDIYSTNIPIRLFVEQFSRQIQEYTIKKGYRPFGTLLLIVGDNKMYRVD<br>PSGSYTSLQVGTIGREYTESGRLLERRKGMLDDNISTCVECIREYCGRSVKSEDIDIGVYRGQEFRVYSKEE<br>VQEVFDSINKI |
| PSA3 | B,P<br>*H1B,H2P* | MSSNTFTEEGRLLQTEYAIKNVSKGGTIIGLVCKDGVILLGINKTELLDEREKIYKINPKVYVSVSGLFGDA<br>MLLKKYGQVKAQDFLYEFDYDCDIDRICNFISEKKQLFTQYNSTRPFGFSFIYAGMKNNKFKLCSTDPSGTI<br>NEWKGVCFGENEDAINNGLRNDFPDEEMDMERGLFEIFKILSKVTECSAKDHKKYEILYFKNEESRFLEFEE<br>IENILLRIEEEKNKK |
| PSA4 | C,Q/<br>*H1C,H2Q* | MEYENALGIFSPDGRLIQVEYAQQASETIIGLVCKDGVILLGINKTELLDPVDKDLNIWYTFSGIKPDSYKV<br>LNEARLICRNYKIKTGTNISFDELAYELSLYKQKFTLDSSMRPFGIRSILLQVKDMAKIYVLEPDGNYSEYK<br>CGAVGQKSVSVCEYLEKCEEEDIIFRSVSGLGTVVQSDKNKVMSHVISKDEIRRVEDETVSQIISTVSVK |
| PSA5 | D,R<br>*H1D,H2R* | MSIVSRQNANTYSAEGRLYQVEYAMQAMNLGTSSIGIKTKDYVLLASEKKIISKLQNPSSVKKHYRVYDHIA<br>LGFSGISADVKTIVDKSRNFAINHEYLYDENCKVERLLEHLADLSLNFDKKEADEKIFSRPFGASLLIIGYD<br>TEPRLFSLDPSGSYLEYHAKAIGSGSEVIENMLEQEFDPNVDINSGLKNILNMLSKVMKDKINNFNVEITAI<br>TKNECKILTPEEIEQFLE |
| PSA6 | E,S<br>*H1E,H2S* | MNSQTDYTNYIIFNPEGKIKQLEFINNTVQLGSTVVALKNKSFGVFVTYNEKRSKFALQQKKIFPINSKSLF<br>SFSGITNDGTKIVKYLKNSTVFENIRKGRDIHPIHVFDDLCYSACIRTLTNGNRLYGVQGLLLTDYQGISLV<br>LFDPKGSAKEVRGMSIGSRSQSCRTILEDECDKFEEYNKEELVRLGIKALRNAYPETGVLNKDNVDIWILET<br>NQESKQIKSEEYLQ |
| PSA7 | F,T<br>*H1F,H2T* | MAILDVDTVYTNTGDILQIGYAQKAADNGNTAICMKNKKGLIMIAEKPIESKLYVSEKNFRIKKVNNSIFQI<br>SSGIETDLVYINENLKNNLISEKHSNDMDVSHESVRNQIVNIIHQFTRYSGVRPIGINLLTCSKYKNEYKIL<br>QTDCTGKSLFFKSSVIGKGSRIVKTELEKLNLENMEIRDLVENGIRILYKSYDPLKDKPFDIEIGIMCEETN<br>GEFVRLEKNQYSEIIEKYKDFSVDGEE |
| **PSB1** | N,2<br>*H1N,H22* | MVAYDNKNNEGFSPNLPEMTG**T**TIMAVKYADGILIGADCRTSMGTYVSSRFTDKLTKISDNIYCCRSGSAAD<br>TQAITQYITELVQRSSFIDKEIPSVKKAAMAAKDIIYRYPNMLAGLIIAGYDTKPRIFNISLGGTMTEAEWQ<br>IGGSGSAYIYGLCDTTFKPNMNLEEALEFVKLAVTCAIKRDNASGGCIRMASITREGVQRFFYSGDKILNST |
| **PSB2** | H,V<br>*H1H,H2V* | MFTKTG**T**TIVGVKYKGGVVICADTRSTSGPIVADKNCSKIHYISDNIQACGAGTSADITRVTRKASKVLSIF<br>SKTYNRLPRVSHCVRTCQLHLHPYQGHISAALVVGGVDDTGAHLYDVYPHGSSNSVSYTALGSGSLAAISIL<br>EAGYKDMNREEAMELACAAIEAGIMNDLYSGSNIDVCVISQEGREMFRNYKKVGVRNEVPKFQYPRSSVKIL<br>KEDIYKYIEEK |
| PSB3 | I,W<br>*H1I,H2W* | MSDISQHYGGSLLAMIGKSSVAFLSDKRLGSGPISVSKNFTKIYSLTPRLFFGFTGLVSDGEMLFKKIRKNY<br>NLFVQDNNKDMEPSELSNMISYILYQKRLQPYYVAVIVCGMTLDKKPYASSMDCIGAMKETSEFVTSGTASK<br>NLMGLSEALFYPEMEDEDLFTTSVQTFLNSSDRDTFGGMGFECLLINPEGYKRREFVGRCD |
| PSB4 | J,X<br>*H1J,H2X* | MESSVALKGNDFVIIGTDSSVKNSYLVLKREEDKFYNINNKVVFTYLGDQGDAFRTSSFINEKLVYEEIQNN<br>VEITPKVTANVIQKTLYDNLRSHPKNCYFLVGGLSQDGPELYSVDLYGSLHENDFMAVGISTYFCYGVLDKE<br>YHKNITKEDGIKIIQKCFDVLKQRCSVDISNIEIKIVSKEGVETINKVL |
| **PSB5** | K,Y<br>*H1K,H2Y* | MEKLFTGDIMEIQNTKVDSAFMKNKIVPYKG**T**TTLAFIFQGGMVIAVDSRASAGSYIASQNVHKVIRVNKHL<br>IGTMAGGASDCYFWEKKMGLYAKLYELKNNKRISVSAASMYLSNCVYSYKGQGLSLGSMVCGYDGDKPVIYY<br>VDDAGQRLSGDLFSVGSGSTIAYGVLNESYRFDLTKEEALNLGKKAIWHATHRDAYSGGNVNLYFMDKNGWE<br>HLGTFDVDKFEQ |
| PSB6 | L,Z<br>*H1L,H2Z* | MFTLQANTKDQIIKDLNIGDLTIRDLNIKDLNIGDSSTVNLFKVDIPLLSNEIPLDFTFDNLPSNKKEIFES<br>FDSFTDFVEGKTKSQESKFNPYEDNSGSTVSIRLNNSIIIAADTRHCSEMGIYSRNTSKIFRIGDFLLTITG<br>FYADGYELYNRLKYQVQIYESFNKISIHSLANLASKIMYSKRLFPYYSYVTLSGFEGDNPYVYSFDCLGHFE<br>EVDSVCNGSGSPLIQPLLDSTIEKKNWAGENYEVTEEYVKDIVRRGFNAASERDVKTGDNVEIWIIKKDGMT<br>KEYERLRQD |
| PSB7 | M,1<br>*H1M,H21* | MKNFVIGSGVISLRYKNGVITCTDTQASYGNLCKFNDVRRIFRLSSNTLISLSGEISDIQFLMNELNKLNES<br>DPVKMSPRGYLNLVQGILYNKRSRVEPLNVSVSIVGVDDDDFLVSCVNHLGNFYEDNIVCTGLSNMIALPFL<br>RTCNVLDLERDEAISLVEKAMTVMCYRSCRSSNRIQIGVVEKGLVDISDPYVLNTDWQVGHNEEEIVL |

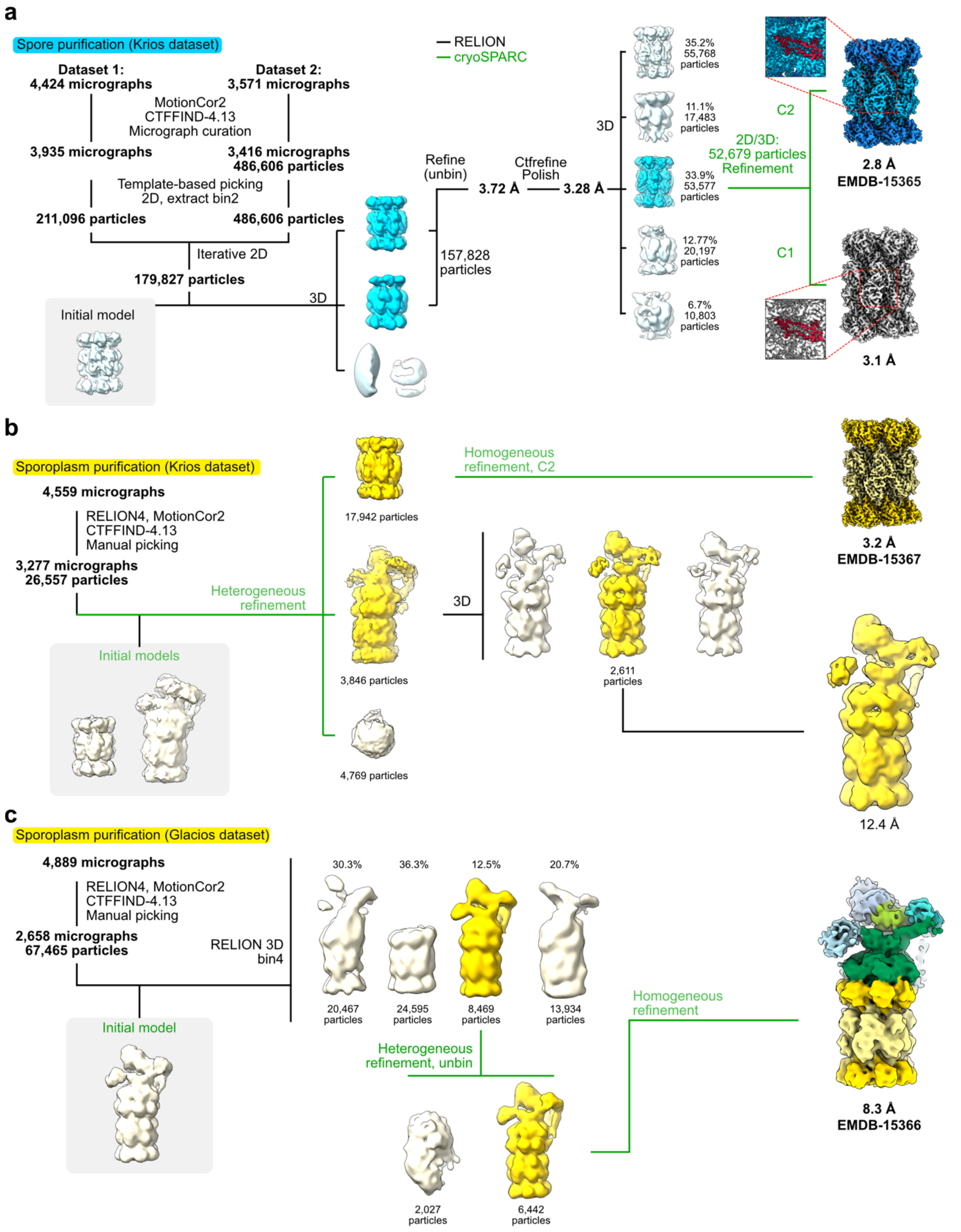

**Supplementary Figure 1. Cryo-EM data processing scheme**. **a-c** Data collection and processing procedures for the two spore-derived datasets, the sporoplasm-derived 20S dataset, and the sporoplasm-derived 26S dataset, respectively. Steps performed in cryoSPARC[1] are represented by green typeface, while steps in Relion[2] are denoted by black typeface.

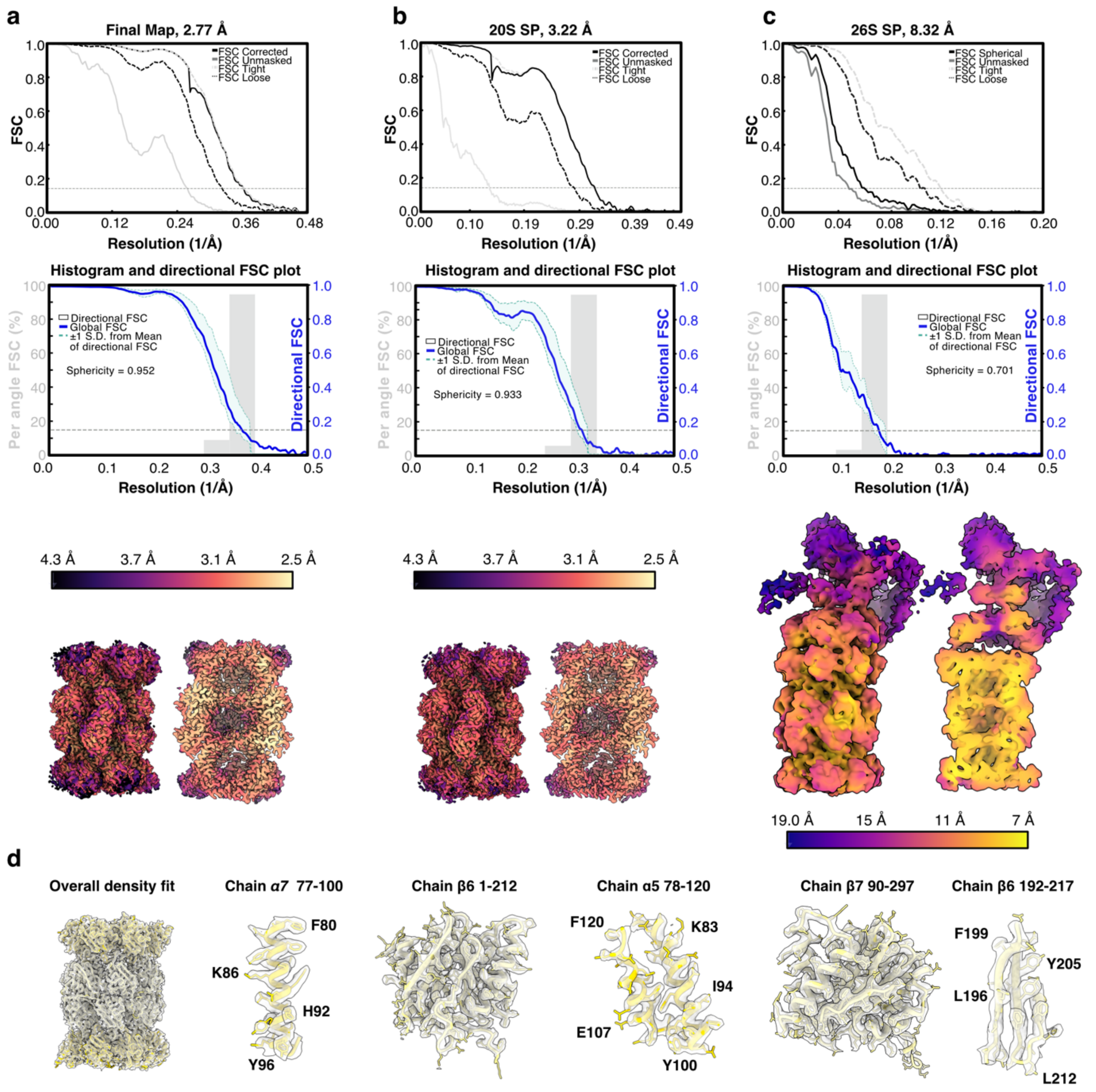

**Supplementary Figure 2**. **Overall and local resolution estimation. a-c** Overall and local resolution estimation of the spore derived 20S (a), the sporoplasm derived 20S (b) and the sporoplasm derived 26S proteasome. (a-c), top to bottom: The cryoSPARC generated Fourier Shell Correlation (FSC) curves are shown above the corresponding 3DFSC plot. The bottom row displays the local resolution surface representations of the respective maps (left) and their corresponding slap view (right) **d** Density and model fit examples of the full spore 20S map, single chains, and selected areas. Residue range and single residues are indicated.

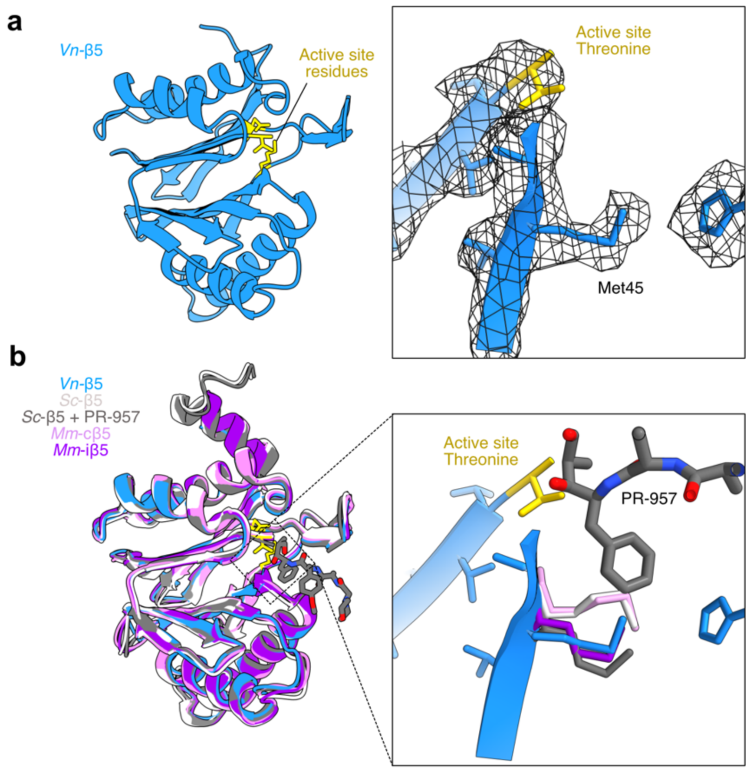

**Supplementary Figure 3**. **Conformational variation of the Met45 residue in β5 subunits. a** Cryo-EM model and electron density mesh map (right) for the *V. necatrix* β5 subunit. Active site residues Thr1, D17, and Lys33 are highlighted in gold, with sidechains shown as sticks. Residues surrounding Met45 are removed for clarity. **b** Superposition of β5 subunits from *V. necatrix* (blue), *S. cerevisiae* (PDB 5CZ4[3], light grey), *S. cerevisiae* in complex with the PR-957 inhibitor (3UN4[4], dark grey), the *Mus musculus* constitutive proteasome (PDB 3UNE[4], pink), and the *M. musculus* immunoproteasome (PDB 3UNH[4], purple). A rotated and magnified superposition of the Met45 residues (right) demonstrates the distinct conformational change to Met45 seen in *S. cerevisiae* upon PR-957 binding. A similar conformation difference is associated with selective binding of the inhibitor to *M. musculus* immunoproteasomes over constitutive proteasomes.
