## Supplementary material for "Reductive evolution in the structure of the microsporidian proteasome": Description of Supplementary Data

**Description of Additional Supplementary Files**

File name: Supplementary Data 1

Description: Mass spectrometry data for proteasome samples derived from dormant spores. As the available *Vairimorpha necatrix* genome is not yet annotated, potential identities for hits were defined via a pblast against yeast proteomes. Proteins with no pblast hit above a cut-off E-value of 0.05 were considered not-identifiable (*N.I.*). Proteasome CP subunits are highlighted in green, and incorrectly identified yeast hits for CP proteins are noted in red.

File Name: Supplementary Data 2

Description: Protein sequences for proteasome subunits in microsporidians with available genome assemblies and for selected outgroups.
